# Biochemical reconstitutions reveal principles of human *γ*-TuRC assembly and function

**DOI:** 10.1101/2020.09.21.306845

**Authors:** Michal Wieczorek, Shih-Chieh Ti, Linas Urnavicius, Kelly R. Molloy, Amol Aher, Brian T. Chait, Tarun M. Kapoor

## Abstract

The formation of cellular microtubule networks is regulated by the *γ*-tubulin ring complex (*γ*-TuRC). This ∼2.3 MDa assembly of >31 proteins includes *γ*-tubulin and GCP2-6, as well as MZT1 and an actin-like protein in a “lumenal bridge”. The challenge of reconstituting the *γ*-TuRC has limited dissections of its assembly and function. Here, we report a complete biochemical reconstitution of the human *γ*-TuRC (*γ*-TuRC-GFP), a ∼35S complex that nucleates microtubules *in vitro*. We extend our approach to generate a stable subcomplex, *γ*-TuRC^mini^-GFP, which lacks MZT1 and actin. Using mutagenesis, we show that *γ*-TuRC^mini^-GFP nucleates microtubules in a guanine nucleotide-dependent manner and proceeds with similar kinetics as reported for native *γ*-TuRCs. Electron microscopy reveals that *γ*-TuRC-GFP resembles the native *γ*-TuRC architecture, while *γ*-TuRC^mini^-GFP adopts a partial cone shape presenting only 8-10 *γ*-tubulin subunits and lacks a well-ordered lumenal bridge. Our structure-function analysis suggests that the lumenal bridge facilitates the self-assembly of regulatory interfaces around a microtubule-nucleating “core” in the *γ*-TuRC.

## Introduction

The *γ*-tubulin ring complex (*γ*-TuRC) is a 2.3 MDa macromolecular assembly that is required for proper microtubule network formation in eukaryotes (Knop et al., 1997; Raff et al., 1993; Stearns and Kirschner, 1994; Zheng et al., 1995). In vertebrates, the *γ*-TuRC contains at least 8 proteins, including the tubulin-like GTPase *γ*-tubulin, *γ*-tubulin ring complex proteins (GCPs) 2-6, and the mitotic-spindle organizing proteins associated with a ring of *γ*-tubulin (MZT1 and MZT2) (Murphy et al., 2001; Teixidó-Travesa et al., 2010; Hutchins et al., 2010). The stoichiometry, location, and structures of these proteins in the context of the native *γ*-TuRC were recently identified using high-resolution cryo-electron microscopy (cryo-EM) (Wieczorek et al., 2019; Liu et al., 2019; Wieczorek et al., 2020; Consolati et al., 2020). These studies showed that the *γ*-TuRC is an asymmetric, cone-shaped structure, in which GCP2-6 orient 14 *γ*-tubulin molecules in a helical arrangement that is poised to nucleate microtubules. Unexpectedly, the *γ*-TuRC also contains a prominent feature inside its conical structure, termed the “lumenal bridge”. The lumenal bridge is composed of two copies of MZT1 that associate with the N-terminal *α*-helical domains (NHDs) of GCP3 and GCP6 in structurally-mimetic “modules”, along with an actin-like protein (Wieczorek et al., 2020). However, the precise role of the lumenal bridge in the assembly and microtubule-nucleating activity of the *γ*-TuRC is currently unclear.

Much of our biochemical understanding of the *γ*-TuRC comes from studies of heterotetramers of GCP2, GCP3, and *γ*-tubulin, also termed the *γ*-tubulin small complex (*γ*-TuSC) (Oegema et al., 1999). Purified recombinant *S. cerevisiae* and *D. melanogaster γ*-TuSCs can nucleate microtubules in bulk assays, but with relatively weak efficiencies compared to native *γ*-TuRCs or multimeric *γ*-TuSC assemblies (Oegema et al., 1999; Vinh et al., 2002; Kollman et al., 2010). *S. cerevisiae γ*-TuSC mutants have been used to demonstrate that GTP binding to *γ*-tubulin is essential for its microtubule nucleating activity (Gombos et al., 2013). However, it is currently unclear if GTP also regulates the function of the *γ*-TuRC holocomplex. Moreover, whether the complete *γ*-TuRC structure is needed for efficient microtubule nucleation, or whether other well-ordered minimal subcomplexes exhibit native *γ*-TuRC-like activity, are open questions. This is largely due to a lack of a reconstitution system with which to study the vertebrate *γ*-TuRC.

Here, we successfully reconstitute the *γ*-TuRC using a set of ten recombinant human proteins: *γ*-tubulin, GCP2, GCP3, GCP4, GCP5, GCP6, NEDD1, MZT1, MZT2, and β-actin. Using this reconstitution strategy, we engineer a subcomplex that lacks the lumenal bridge components MZT1 and *β*-actin. We use electron microscopy to show that this subcomplex adopts a semi-conical shape organizing ∼8 copies of *γ*-tubulin into an arrangement that mimics the native *γ*-TuRC, but which lacks an ordered “overlap” region and lumenal bridge. Despite its partial structure, this subcomplex can nucleate individual microtubules in a manner analogous to native *γ*-TuRCs (Consolati et al., 2020). We further validate our functional assays by introducing a mutation into *γ*-tubulin that abrogates the subcomplex’s microtubule-nucleating activity without interfering with its overall assembly. Our results show that the human *γ*-TuRC can be reconstituted using a limited set of purified components, and suggest that the role of the lumenal bridge is to build regulatory interfaces around a microtubule nucleation-competent “core”.

## Results

### Reconstitution of a stable *γ*-TuRC-sized complex from ten recombinant human proteins co-overexpressed in insect cells

To address the need for a biochemically tractable system with which to study the function and assembly of the vertebrate *γ*-TuRC, we asked whether the human complex could be reconstituted with recombinant proteins. We tested several strategies, including individually expressing *γ*-TuRC components either in human cells under cytomegalovirus (CMV) promoters or in insect cells using the baculovirus expression system. Encouraging yields were only obtained when a stoichiometric excess of *γ*-tubulin, GCP2, and GCP3 were co-overexpressed with GCP4, GCP5, GCP6, NEDD1, MZT1, *β*-actin, and ZZ-tagged MZT2A (hereafter, MZT2; (Wieczorek et al., 2020)) in insect cells infected with polycistronic baculoviruses (Figure 1A; see Methods). Recently, we reported that the structured portion of MZT2 (a.a. 35-80) adopts a MZT1-like fold and binds to the flexible N-terminus of GCP2 (GCP2-NHD) (Wieczorek et al., 2020). Guided by these structural data, mEGFP was fused to MZT2’s unstructured C-terminus (a.a. 81-158). We collectively refer to this set of ten recombinant proteins as “*γ*-TuRC-GFP”. Gratifyingly, and as detailed further below, a stable complex containing all *γ*-TuRC-GFP proteins could be isolated from infected insect cell lysates using three purification steps (Figure 1A): 1) IgG affinity capture and tobacco etch virus (TEV) protease release; 2) preparative gel filtration (Figure 1B-C); and 3) sucrose density gradient centrifugation (Figure 1C).

**Figure 1.**
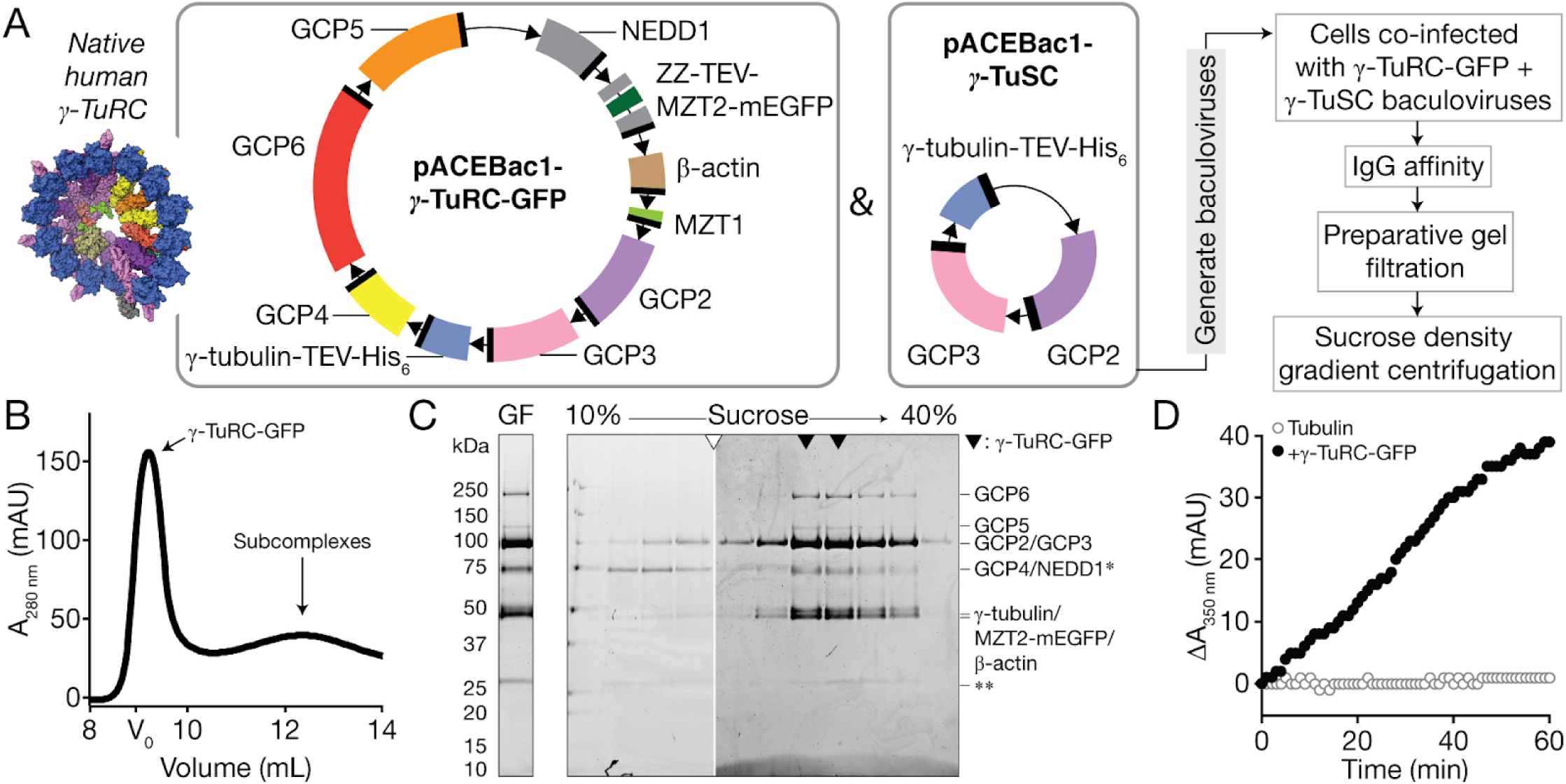
Reconstitution, purification, and functional characterization of recombinant *γ*-TuRC-GFP. A) Strategy for expressing and purifying *γ*-TuRC-GFP. Left, inset: Surface representation of a model for the native human *γ*-TuRC (PDB ID: 6V6S, 6×0U, and 6×0V (Wieczorek et al., 2019, 2020)). Middle: Schematics of recombinant protein constructs used to express *γ*-TuRC-GFP in insect cells. Donor plasmids containing 10 *γ*-TuRC-GFP proteins (pACEBac1-*γ*-TuRC-GFP) and *γ*-tubulin-TEV-His_6_/GCP2/GCP3 (pACEBac1-*γ*-TuSC) used to generate bacmids are indicated. “ZZ” = IgG-binding purification tag; TEV = tobacco etch virus protease site. Coding fragments are colored according to the convention in (Wieczorek et al., 2019) and as in the inset model. Right: *γ*-TuRC-GFP expression and purification scheme. B) Superose 6 Increase elution profile of *γ*-TuRC-GFP IgG eluate. Fractions containing intact *γ*-TuRC-GFP are indicated. The void volume of the column was estimated by gel filtering Dextran Blue 2000 in gel filtration buffer. C) Coomassie-stained SDS-PAGE analysis of *γ*-TuRC-GFP after gel filtration (GF; left) and sucrose gradient centrifugation (right). Hollow triangle indicates where gels were cropped. Solid triangles indicate peak *γ*-TuRC-GFP fractions. Single asterisk indicates a ∼70 kDa contaminant that does not co-sediment with *γ*-TuRC-GFP components. Double asterisk indicates a contaminant at ∼25 kDa likely to be IgG light chain (Murphy et al., 2001). MZT1 (∼8 kDa) is not visible on the gel but is detected in LC-MS/MS (Figure S1B). D) Turbidity-based microtubule nucleation assay of reactions containing tubulin (20 μM) and GTP (1 mM) alone (hollow grey circles) or in the presence of 0.25 nM *γ*-TuRC-GFP (solid black circles).

We characterized the biochemical integrity of purified *γ*-TuRC-GFP in two ways. First, the sucrose density gradient used as a purification step was calibrated with standards of known S values (Figure 1C and Figure S1A), which indicated that *γ*-TuRC-GFP sediments to 32-38 S, a range that is larger than individual *γ*-TuSCs (9.8 S) but comparable to the native *D. melanogaster γ*-TuRC (35.5 S; (Oegema et al., 1999)). The purified complex was also analyzed by mass spectrometry (Figure S1B), which confirmed the presence of all ten overexpressed *γ*-TuRC-GFP proteins listed above. Notably, the sucrose gradient removed a contaminant at ∼70 kDa likely corresponding to insect cell Hsp70 (Figure 1C).

To assess the activity of the purified complex, we incubated *γ*-TuRC-GFP (∼0.25 nM; estimated using recombinant human *γ*-tubulin as a standard in quantitative western blots; see Methods) with *α*/*β*-tubulin (20 μM) and GTP (1 mM), and measured the change in optical density, or “turbidity” (Gaskin et al., 1974), of the reaction over time. An increase in turbidity interpreted as the formation of microtubule polymer was observed only in the presence of both *γ*-TuRC-GFP and tubulin (Figure 1D); no obvious increase in turbidity was detected in tubulin-(Figure 1D) or *γ*-TuRC-GFP-only (data not shown) controls. Together, our results indicate that a functional *γ*-TuRC-sized complex can be reconstituted with a set of ten recombinant human proteins expressed in insect cells.

### Reconstitution of a stable subcomplex lacking MZT1 and *β*-actin

We next focused on examining the role of the lumenal bridge, which spans across the inside of the *γ*-TuRC cone and has been proposed to be essential for both the assembly and function of the native complex (Figure 2A; (Liu et al., 2019)). We generated a polycistronic bacmid containing all *γ*-TuRC-GFP proteins except for MZT1 and *β*-actin (Figure 2B). We collectively refer to the resulting set of eight proteins as “*γ*-TuRC^mini^-GFP”. *γ*-TuRC^mini^-GFP proteins were co-overexpressed with a stoichiometric excess of *γ*-tubulin, GCP2, and GCP3 in insect cells and the resulting complex was isolated using the same 3-step purification procedure as for *γ*-TuRC-GFP (Figure 1A). The majority of *γ*-TuRC^mini^-GFP proteins from peak preparative gel filtration fractions (Figure 2C-D) cosedimented together in sucrose density gradients (Figure 2D), albeit to lower estimated S values when compared with *γ*-TuRC-GFP (∼32 S, Figure S1C vs. ∼35 S, Figure S1A). This suggests that *γ*-TuRC^mini^-GFP has a smaller mass than both *γ*-TuRC-GFP and native *γ*-TuRCs (Figure S1A; (Oegema et al., 1999)). Mass spectrometry also confirmed the presence of all 8 *γ*-TuRC^mini^-GFP proteins in the peak sucrose density gradient fractions (Figure S1D). Together, these observations suggest that a complex containing 8 *γ*-TuRC proteins can assemble in the absence of the lumenal bridge components MZT1 and *β*-actin.

**Figure 2.**
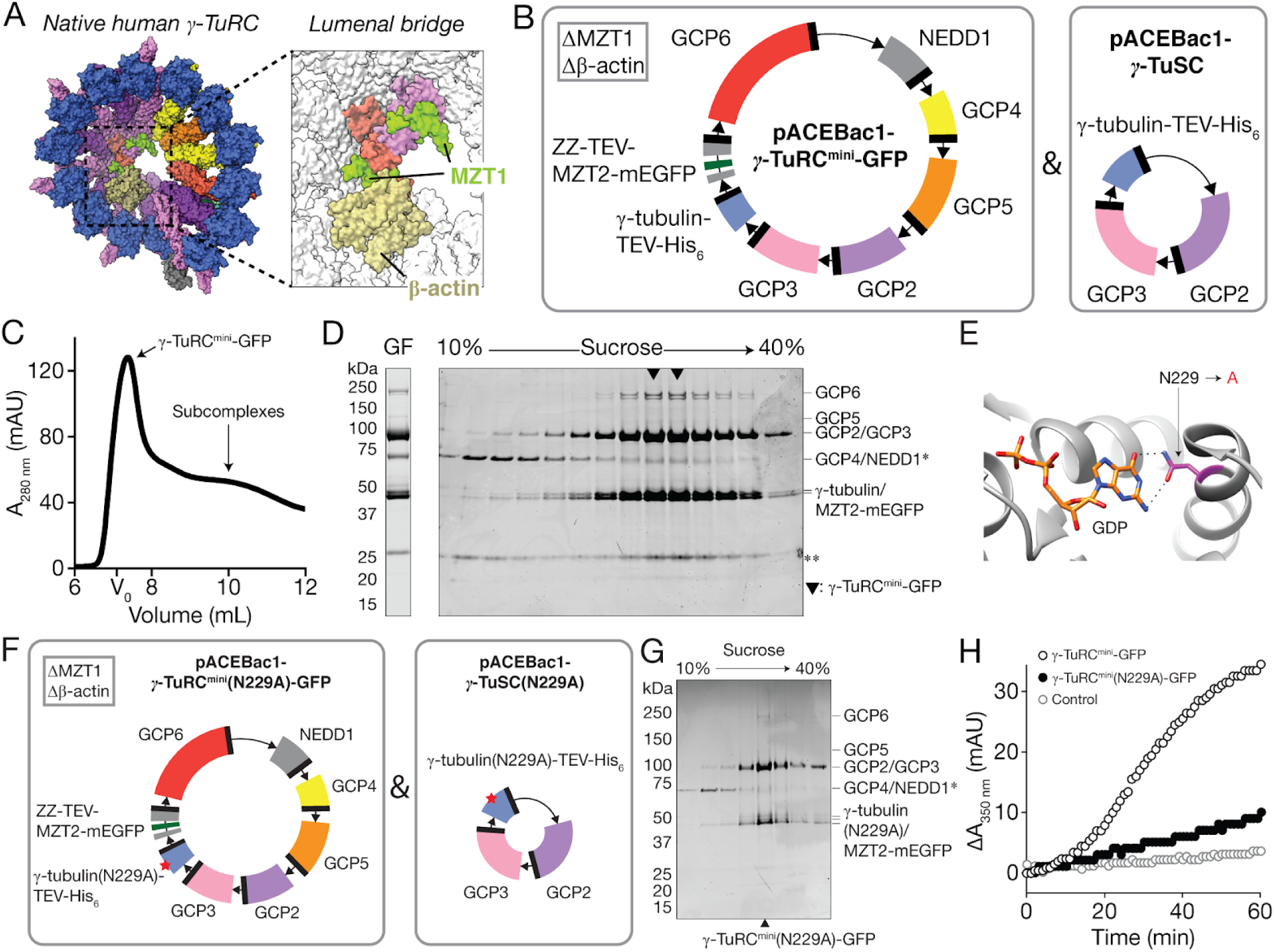
Reconstitution, purification and functional characterization of *γ*-TuRC^mini^-GFP. A) Surface representation of a model for the native human *γ*-TuRC, as in Figure 1A. Inset shows the lumenal bridge and highlights models for *β*-actin and MZT1. B) Schematics of recombinant protein constructs used to express *γ*-TuRC^mini^-GFP in insect cells. Donor plasmids containing 8 *γ*-TuRC^mini^-GFP proteins (pACEBac1-*γ*-TuRC^mini^-GFP) and *γ*-tubulin-TEV-His_6_/GCP2/GCP3 (pACEBac1-*γ*-TuSC) used to generate bacmids are indicated. C) Superose 6 elution profile of *γ*-TuRC^mini^-GFP IgG eluate. Fractions containing intact *γ*-TuRC^mini^-GFP are indicated. The void volume of the column was estimated by gel filtering Dextran Blue 2000 in gel filtration buffer. D) Coomassie-stained SDS-PAGE analysis of *γ*-TuRC^mini^-GFP after gel filtration (GF; left) and sucrose gradient centrifugation (right). Solid triangles indicate peak *γ*-TuRC^mini^-GFP fractions. Single asterisk indicates a ∼70 kDa contaminant that does not co-sediment with *γ*-TuRC-GFP components. Double asterisk indicates a contaminant at ∼25 kDa likely to be IgG light chain (Murphy et al., 2001). E) View of the nucleotide-binding site in the X-ray crystal structure of GDP-bound human *γ*-tubulin (PDB ID: 3CB2; grey cartoon representation). Residue N229 (magenta), which was mutated to alanine in this study, and GDP (colored by element) are indicated. Predicted hydrogen bonds formed between N229 and the guanine moiety of GDP are shown (dotted black lines). F) Schematics of the recombinant protein constructs used to express *γ*-TuRC^mini^(N229A)-GFP in insect cells. Donor plasmids containing the 8 *γ*-TuRC^mini^(N229A)-GFP proteins (pACEBac1-*γ*-TuRC^mini^(N229A)-GFP; left) and *γ*-tubulin(N229A)-TEV-His_6_/GCP2/GCP3 (pACEBac1-*γ*-TuSC(N229A); right) are indicated. The N229A point mutation is highlighted by a red star in the *γ*-tubulin(N229A) coding fragments. ZZ = IgG-binding purification tag; TEV = tobacco etch virus protease site; coding fragments are colored as in Figure 1A. G) Coomassie-stained SDS-PAGE analysis of *γ*-TuRC^mini^(N229A)-GFP sucrose gradient centrifugation fractions. Asterisk indicates a ∼70 kDa contaminant that does not co-sediment with *γ*-TuRC^mini^(N229A)-GFP components in the sucrose gradient. Peak *γ*-TuRC^mini^(N229A)-GFP fraction is indicated by a solid triangle on the bottom of the gel. H) Turbidity-based microtubule nucleation assay of reactions containing tubulin (20 μM) and GTP (1 mM) alone (hollow grey circles), in the presence of 0.25 nM *γ*-TuRC^mini^-GFP (hollow black circles), or in the presence of 0.25 nM *γ*-TuRC^mini^(N229A)-GFP (solid black circles).

We next asked whether *γ*-TuRC^mini^-GFP is capable of nucleating microtubules, despite lacking MZT1 and *β*-actin. As a control, we also introduced a point mutation in the nucleotide-binding site of *γ*-tubulin (N229A; Figure 2E). Though the precise role of GTP hydrolysis in *γ*-TuRC function is not known, this mutation is predicted to significantly reduce human *γ*-tubulin’s nucleotide affinity and microtubule-nucleating activity based on previous bulk nucleation assays with yeast *γ*-TuSC assemblies harboring a homologous *γ*-tubulin mutation (Gombos et al., 2013). We co-overexpressed *γ*-tubulin(N229A) along with the other 7 *γ*-TuRC^mini^-GFP components (Figure 2F) and purified the resulting subcomplex, *γ*-TuRC^mini^(N229A)-GFP, using a similar strategy as for *γ*-TuRC-GFP (Figure 1A). *γ*-TuRC^mini^(N229A)-GFP sedimented to a qualitatively similar position in sucrose gradients as *γ*-TuRC^mini^-GFP (Figure 2G) and contained all 8 overexpressed proteins as judged by mass spectrometry (Figure S1E), which suggests that the N229A mutation does not interfere with the assembly of *γ*-TuRC^mini^-GFP. Incubation of *γ*-TuRC^mini^(N229A)-GFP (∼0.25 nM) with tubulin (20 µM) and GTP (1 mM) did not produce a substantial increase in signal in the turbidity assay (Figure 2H). Under the same conditions, however, the presence of “wild-type” *γ*-TuRC^mini^-GFP did produce an increase in turbidity in a manner that we note is qualitatively similar as *γ*-TuRC-GFP (Figure 2H and Figure 1D). Together, these data indicate that *γ*-TuRC^mini^-GFP can nucleate microtubules despite lacking MZT1 and *β*-actin, key structural components of the native *γ*-TuRC lumenal bridge.

### *γ*-TuRC^mini^-GFP nucleates individual microtubules with similar kinetics as the native *γ*-TuRC

To gain insight into the kinetics of microtubule nucleation from *γ*-TuRC^mini^-GFP, we immobilized the complex to a surface and imaged individual microtubule nucleation events using single molecule total internal reflection fluorescence (TIRF) microscopy (Figure 3A). Briefly, *γ*-TuRC^mini^-GFP was adhered to glass coverslips functionalized with a biotinylated GFP nanobody (Fridy et al., 2014), and a mixture containing fluorescent tubulin (15 μM and ∼5% X-rhodamine:tubulin labeling ratio) and GTP (1 mM) was introduced to the flow chamber. Within minutes, the polymerization of individual microtubules on the coverslip surface could be observed in the tubulin channel; in coverslips prepared without immobilized *γ*-TuRC^mini^-GFP, microtubule polymerization was not observed during the time course of the experiment (typically 20-30 minutes; Figure 3B). ∼95% of microtubule nucleation events were observed to initiate from *γ*-TuRC^mini^-GFP puncta (Figure S1F). The growth rates and catastrophe frequencies of microtubules nucleated by *γ*-TuRC^mini^-GFP matched published values for those of plus ends grown off of GMPCPP seeds (Brouhard et al., 2008; Gardner et al., 2011), axonemes (Walker et al., 1988), and native *γ*-TuRCs (Consolati et al., 2020; Thawani et al., 2020) (Figure S1G-H). Moreover, the position of fluorescent-tubulin “speckles” along the growing microtubule’s length did not change relative to the position of microtubule end-bound complexes (Figure S1I-K; (Waterman-Storer et al., 1998)), indicating that *γ*-TuRC^mini^-GFP nucleates microtubules that grow from their distal plus ends.

**Figure 3.**
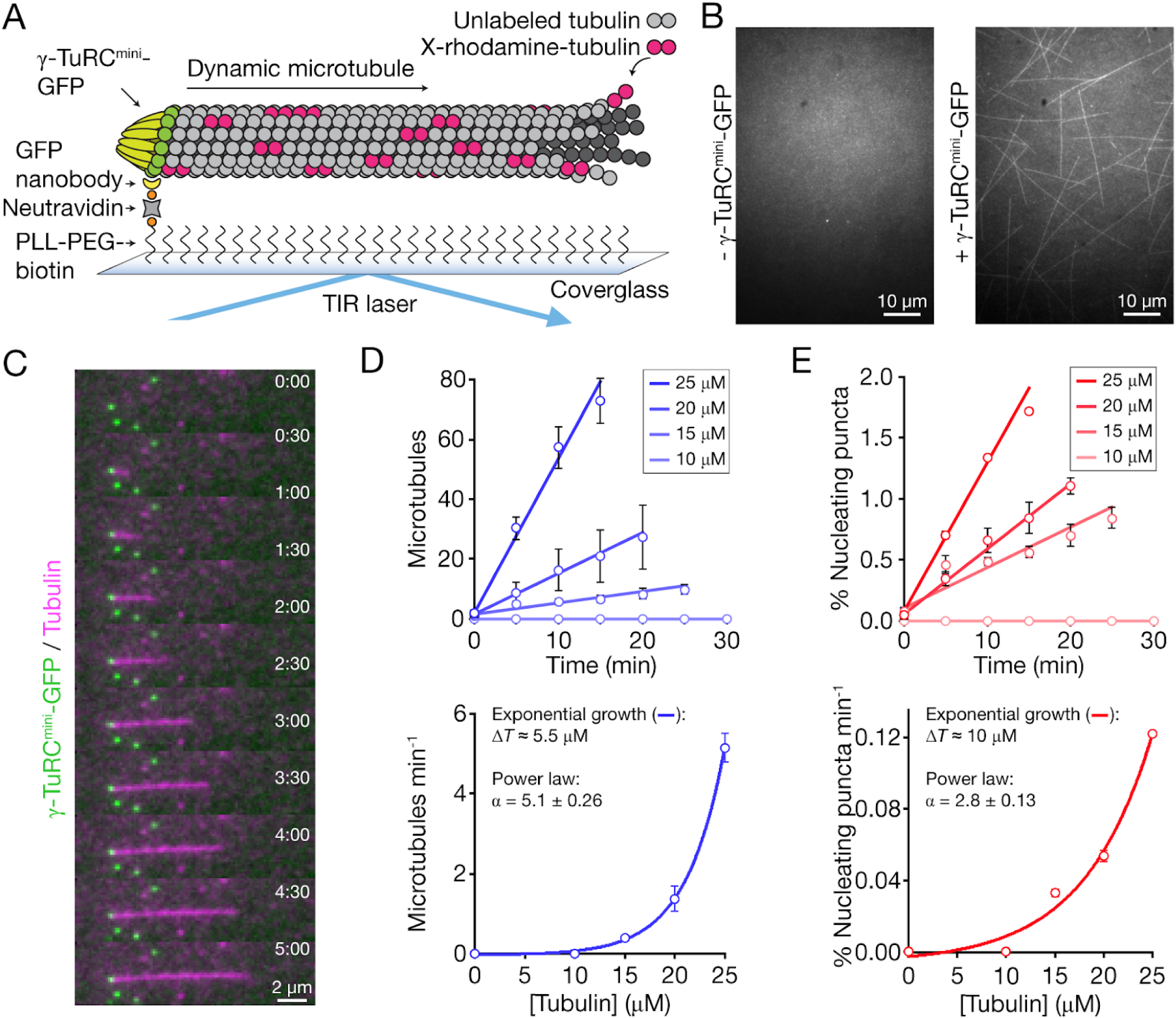
*γ*-TuRC^mini^-GFP nucleates individual microtubules with native *γ*-TuRC-like kinetics. A) Schematic of the TIRF-based assay developed to visualize individual microtubule nucleation events from *γ*-TuRC^mini^-GFP. B) Fields of view displaying the tubulin channel after 20 min in the absence (left) or presence (right) of surface-immobilized *γ*-TuRC^mini^-GFP. C) Two-color overlay time lapse sequence of a microtubule nucleated by and stably attached to a *γ*-TuRC^mini^-GFP punctum. Time is reported as minutes:seconds. D) Top: Plot of the cumulative number of microtubules nucleated over time at increasing tubulin concentrations. Data were fitted using weighted linear regression (solid lines). Bottom: Plot of the rate of microtubule formation over time as a function of tubulin concentration. Data were fitted to an exponential growth model of the form y = y_0_ + Aexp(x/t), where x = [Tubulin] in µM, and ΔT = t/ln(2) is an estimate of the change in tubulin concentration (in µM) needed to double the nucleation rate. The non-zero nucleation rates were also fitted to a power law of the form ln(y) = *α*ln(x) + ln(*β*); the value for the exponent *α* is indicated. E) Top: Plot of the cumulative percentage of *γ*-TuRC^mini^-GFP puncta that nucleated a microtubule over time at increasing tubulin concentrations. Data were fitted using weighted linear regression (solid lines). Bottom: Plot of the rate at which the number of *γ*-TuRC^mini^-GFP-nucleating puncta increases over time as a function of tubulin concentration. Data were fitted to an exponential growth model of the form y = y_0_ + Aexp(x/t), where x = [Tubulin] in µM, and ΔT = t/ln(2) is an estimate of the change in tubulin concentration (in µM) needed to double the number of *γ*-TuRC^mini^-GFP-nucleating puncta. The non-zero nucleation rates were also fitted to a power law of the form ln(y) = *α*ln(x) + ln(*β*); the value for the exponent *α* is indicated. Experiments shown in D) - E) were repeated 2-3 times, and all experiments contained at least 500 *γ*-TuRC^mini^-GFP puncta per field of view. Error bars represent s.e.m. Errors were propagated manually.

The microtubule nucleating activity of the native human *γ*-TuRC has been found to be surprisingly low (<1% nucleation efficiency; (Consolati et al., 2020)). We also found that many *γ*-TuRC^mini^-GFPs (>95%) did not nucleate microtubules during the time course of our experiments. To further characterize the microtubule-nucleating efficiency of *γ*-TuRC^mini^-GFP, we repeated the single-molecule nucleation assay at increasing tubulin concentrations, normalizing for the *γ*-TuRC^mini^-GFP surface density (see Methods). The number of *γ*-TuRC^mini^-GFP-mediated microtubule nucleation events (Figure 3D) and the percentage of *γ*-TuRC^mini^-GFP puncta that nucleated microtubules (Figure 3E) were then measured. Both of these quantities were found to increase linearly over time (Figure 3D-E, upper panels), suggesting that the majority of surface-bound *γ*-TuRC^mini^-GFP molecules are capable of nucleating microtubules, given enough time. The rates of both quantities were plotted and were observed to increase exponentially with tubulin concentration (Figure 3D-E, lower panels), indicating that nucleation from *γ*-TuRC^mini^-GFP is a cooperative process. An exponential fit to the *γ*-TuRC^mini^-GFP-mediated microtubule nucleation rates revealed a characteristic doubling concentration ΔT = ∼5.5 µM tubulin (Figure 3D); similarly, the percentage of *γ*-TuRC^mini^-GFPs that will nucleate a microtubule will double when ΔT = ∼10 µM tubulin (Figure 3E). Consistent with recent reports using native *γ*-TuRCs (Consolati et al., 2020), >10 µM tubulin was required to observe microtubule nucleation from *γ*-TuRC^mini^-GFP, a concentration which is larger than that needed to nucleate from pre-formed microtubule seeds (∼7 µM), but less than that required for spontaneous microtubule assembly (∼20 µM) (Wieczorek et al., 2015). Moreover, a power law fit to the microtubule nucleation rate data in Figure 3D returned an exponent of *α* = 5.1 +/- 0.26, which is remarkably similar to the value of 6.7 reported recently in a similar analysis of the native human *γ*-TuRC (Consolati et al., 2020). Thus, our results suggest that immobilized *γ*-TuRC^mini^-GFP nucleates the plus end growth of single microtubules in a manner that is remarkably similar to the native complex.

### *γ*-TuRC^mini^-GFP adopts a partial ring structure

To determine whether the absence of MZT1 and *β*-actin impacts the architecture of the *γ*-TuRC, we characterized the structures of our reconstituted complexes using negative stain electron microscopy (EM) (see Methods). First, we examined *γ*-TuRC-GFP, which appears as ∼25-30 nm-wide lockwasher-shaped assemblies in raw EM micrographs (Figure 4A), similar to previous analyses of native human *γ*-TuRCs (Murphy et al., 2001). Reference-free 2D classification averages indicated that *γ*-TuRC-GFP adopts a ring-shaped assembly, in which multiple ∼5 nm-wide globular densities laterally associate and are supported by stalk-like structures (Figure 4B). A 3D reconstruction of the EM data revealed a cone-shaped structure with ∼14 globular domains (FSC_0.5_ = ∼30 Å; Figure 4C and Figure S2A-B), which is similar in overall architecture as recent cryo-EM reconstructions of the native human *γ*-TuRC (Wieczorek et al., 2019; Consolati et al., 2020; Wieczorek et al., 2020). Correspondingly, a model of the native complex could be rigid-body fitted into the *γ*-TuRC-GFP density (PDB ID: 6V6S, 6×0U, and 6×0V (Wieczorek et al., 2019, 2020); Figure S2C). A high local correlation score between the fitted model and the *γ*-TuRC-GFP density (Figure S2D) indicates that the asymmetric features (e.g. a lumenal bridge and a helical deviation in the *γ*-tubulin ring; Figure 4C) in the native complex are also present in *γ*-TuRC-GFP (Wieczorek et al., 2019). Together, these data show that purified *γ*-TuRC-GFP stably assembles into an asymmetric, cone-shaped structure, which at the resolution of our negative stain EM data is indistinguishable from the architecture of the native human *γ*-TuRC.

**Figure 4.**
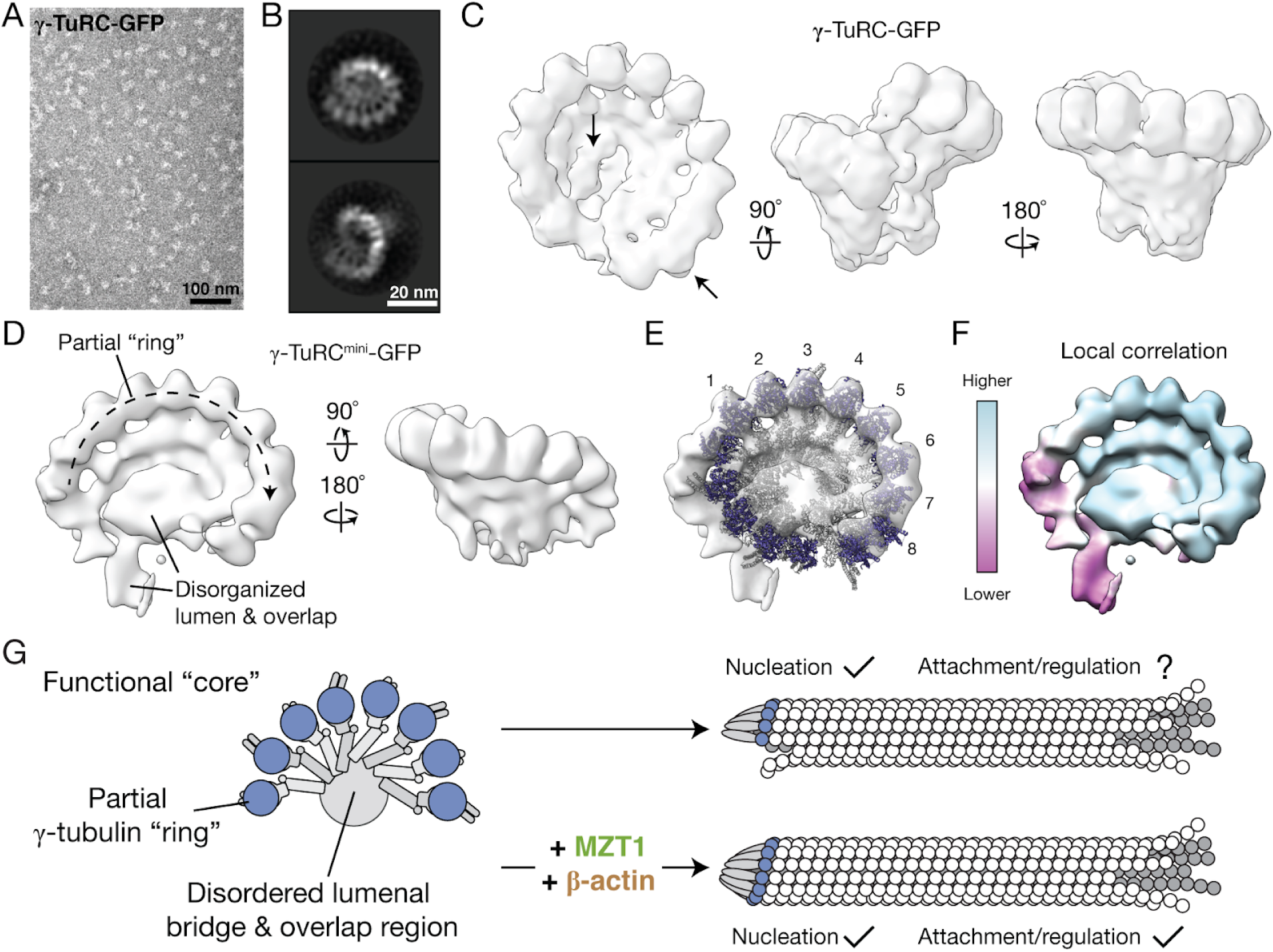
Reconstituted *γ*-TuRC^mini^-GFP adopts a partial, cone-shaped structure. A) Transmission EM micrograph of negatively-stained *γ*-TuRC-GFP. B) 2D class averages showing two orientations of *γ*-TuRC-GFP particles. C) Three views of a 3D reconstruction of *γ*-TuRC-GFP from negative stain EM data. Arrows point to the well-ordered lumenal bridge and overlap regions. D) Two views of a 3D reconstruction of *γ*-TuRC^mini^-GFP from negative stain EM data. E) Rigid-body fit of the native human *γ*-TuRC model from Figure 1A into the *γ*-TuRC^mini^-GFP density map. Globular densities in the partial ring are numbered. F) The *γ*-TuRC^mini^-GFP density map colored according to local correlation of the model fit in E) (see Methods). G) Proposed model for the role of the lumenal bridge in *γ*-TuRC assembly, microtubule nucleation, and downstream regulation.

We next examined the structure of *γ*-TuRC^mini^-GFP. Negative stained *γ*-TuRC^mini^-GFP particles in raw EM micrographs looked qualitatively similar to those from grids containing *γ*-TuRC-GFP (Figure S3A vs Figure 4A). Remarkably, however, 3D classification of >900,000 particles revealed that *γ*-TuRC^mini^-GFP mainly adopts distinct, semi-conical structures typically displaying 8-10 *γ*-tubulin-like globular domains (Figure S3B-C). Native-like *γ*-TuRC structures were not observed in these averages, which is in contrast to *γ*-TuRC-GFP (Figure 4C; data not shown). We generated a 3D reconstruction of a subset of the *γ*-TuRC^mini^-GFP particles that gave the highest resolutions in 3D classifications (FSC_0.5_ = ∼25 Å; Figure 4D and Figure S3B-E; see Methods). The resulting density map revealed that, when viewed from the top, *γ*-TuRC^mini^-GFP contains ∼8 well-defined *γ*-tubulin-like subunits lining the rim of a semi-conical structure with qualitatively similar helical parameters as *γ*-TuRC-GFP (Figure 4D, right panel vs. Figure FC, right panel). Consistent with this observation, a local correlation analysis of a rigid-body fitted model of the native human *γ*-TuRC into the *γ*-TuRC^mini^-GFP density map indicates that *γ*-TuRC^mini^-GFP closely resembles a portion of the native *γ*-TuRC corresponding to ∼8 GCP/*γ*-tubulin subunits (PDB ID: 6V6S, 6×0U, and 6×0V (Wieczorek et al., 2019, 2020); Figure 4E-F). The *γ*-TuRC^mini^-GFP density map also displays additional, less well-resolved *γ*-tubulin-like subunits that appear to deviate from the ring at the edges of the semi-conical structure (Figure 4D). We performed a similar negative stain EM analysis for *γ*-TuRC^mini^(N229A)-GFP, which revealed that *γ*-TuRC^mini^(N229A)-GFP also adopts a partial “ring”-like structure containing ∼8 *γ*-tubulin-like domains (FSC_0.5_ = ∼40 Å; Figure S3F-I). Notably, both *γ*-TuRC^mini^-GFP and *γ*-TuRC^mini^(N229A)-GFP reconstructions lack well-organized lumenal bridge and “overlap” regions (positions 1-2 and 13-14 in the native *γ*-TuRC; (Wieczorek et al., 2019; Liu et al., 2019; Consolati et al., 2020; Wieczorek et al., 2020)) (Figure 4D vs Figure 4C). These results suggest that *γ*-TuRC^mini^-GFP adopts a semi-conical structure that organizes ∼8 *γ*-tubulins into a partial ring that is native *γ*-TuRC-like, yet lacks the lumenal bridge observed in *γ*-TuRC-GFP.

## Discussion

In this study, we have reconstituted the human *γ*-TuRC holocomplex (*γ*-TuRC-GFP) from a set of 10 recombinant proteins co-overexpressed in insect cells. We also reconstitute a subcomplex, *γ*-TuRC^mini^-GFP, and find that it nucleates single microtubules with a similar efficiency as determined for the native human *γ*-TuRC (Consolati et al., 2020). In contrast to *γ*-TuRC-GFP, which assembles into a native-like complex, *γ*-TuRC^mini^-GFP adopts a semi-conical structure that retains some aspects of native *γ*-TuRC architecture (e.g., a partial ring of *γ*-tubulin subunits); however, *γ*-TuRC^mini^-GFP lacks the lumenal bridge.

Previous reconstitution work has focused on the *γ*-TuSC (Gunawardane et al., 2000; Vinh et al., 2002; Lin et al., 2016; Leong et al., 2019), a stable and evolutionarily-conserved subcomplex containing a Y-shaped arrangement of GCP2, GCP3, and two copies of *γ*-tubulin (Kollman et al., 2008). In the presence of the accessory factor Spc110, purified *S. cerevisiae γ*-TuSCs can spontaneously self-assemble into helical assemblies (Kollman et al., 2010). Reconstituted *C. albicans* and *S. pombe γ*-TuSCs display analogous properties, though their self-assembly depends on both Spc110 homologs and the microprotein MZT1 (Lin et al., 2016; Leong et al., 2019). In contrast, our work identifies *γ*-TuRC^mini^-GFP as a stable, partial ring-like subcomplex containing both *γ*-TuSC proteins and the evolutionarily-divergent GCP4, GCP5, and GCP6. The limited resolution provided by negative staining precludes the assignment of GCP subunits in the *γ*-TuRC^mini^-GFP density map (Figure 4D), and our efforts to determine the structure of *γ*-TuRC^mini^-GFP by single particle cryo-EM have not yet been successful (data not shown). Nevertheless, we envision two possibilities: 1) *γ*-TuRC^mini^-GFP polymerizes into an array of 4-5 *γ*-TuSCs; or 2) a recently described stable GCP4/GCP5/GCP4/GCP6-containing subcomplex acts as a polymerization template, with which multiple *γ*-TuSCs can transiently associate (Haren et al., 2020). Previously, we identified extensive electrostatic interactions between the N-terminal GCP6 “belt” (a.a. 130–195) and the lumenal face of at least two *γ*-TuSCs in the native *γ*-TuRC (Wieczorek et al., 2020). The GCP6 belt could conceivably stabilize the assembly of a finite number of *γ*-TuSC subunits with a stable GCP4/GCP5/GCP4/GCP6 *γ*-TuSC-like dimer (Haren et al., 2020).

The finding that *γ*-TuRC^mini^-GFP lacks the lumenal bridge and cannot complete the cone-shaped *γ*-TuRC structure has important implications for the regulation of microtubule nucleation in cells. *γ*-TuRC^mini^-GFP lacks protein-binding interfaces found in the native *γ*-TuRC. One prominent missing interface is the overlap region, which is proximal to the binding site of the CM1 motif of CDK5Rap2 (Wieczorek et al., 2020), a *γ*-TuRC attachment factor and proposed “activator” of the native complex’s microtubule-nucleating function (Choi et al., 2010). Our data raise the possibility that *γ*-TuRC^mini^-GFP-like subcomplexes that can retain robust microtubule-nucleating activity may still form in cells (Figure 4G). Consistent with this idea, depletion of MZT1 in human cells does not eliminate the formation of *γ*-TuRC-like complexes (Cota et al., 2017; Lin et al., 2016), but does abolish ectopic stimulation of microtubule nucleation by CDK5Rap2 overexpression (Lin et al., 2016). Together, our findings reveal the importance of an evolutionarily divergent structural feature - the lumenal bridge - in regulating the assembly of vertebrate *γ*-TuRCs. It is currently unclear whether - and how - *γ*-TuRCs might be biochemically activated into efficient microtubule nucleation templates by associated factors or post-translational modifications such as *γ*-tubulin phosphorylation, as proposed in current models (Kollman et al., 2011). Our reconstitution strategy establishes minimal systems for testing these models using biochemical approaches.

## Acknowledgements

The authors acknowledge B. Graczyk for support in generating DNA constructs used in this study, Drs. J. Luders, M. Rout, and G. Alushin for the generous gift of plasmids, and Dr. S. Liu for access to a plasma cleaner. The authors also thank Dr. H. A. Pasolli, Dr. N. Soplop, M. Ebrahim, J. Sotiris, and H. Ng for fantastic electron microscopy support, and the Rockefeller University Proteomics Resource Center for additional mass spectrometry support. This work was funded by an NIH grant to T.M.K. (R35 GM130234). M.W. is supported by an HFSP fellowship (LT000025/18-L1). L.U. is supported by the Rockefeller University’s Pels Center for Biochemistry and Structural Biology.

## Author contributions

M.W., S.-C.T. and T.M.K. conceived the experiments. M.W., S.-C.T. and A.A. designed constructs. M.W. and S.-C.T. purified proteins. K.R.M. and B.T.C. performed and analyzed mass spectrometry experiments. M.W. and L.U. performed and analyzed electron microscopy experiments. M.W. performed and analyzed functional assays. M.W., S.-C.T., L.U., A.A. and T.M.K. prepared the manuscript.

## Supplementary Information

**Figure S1.**
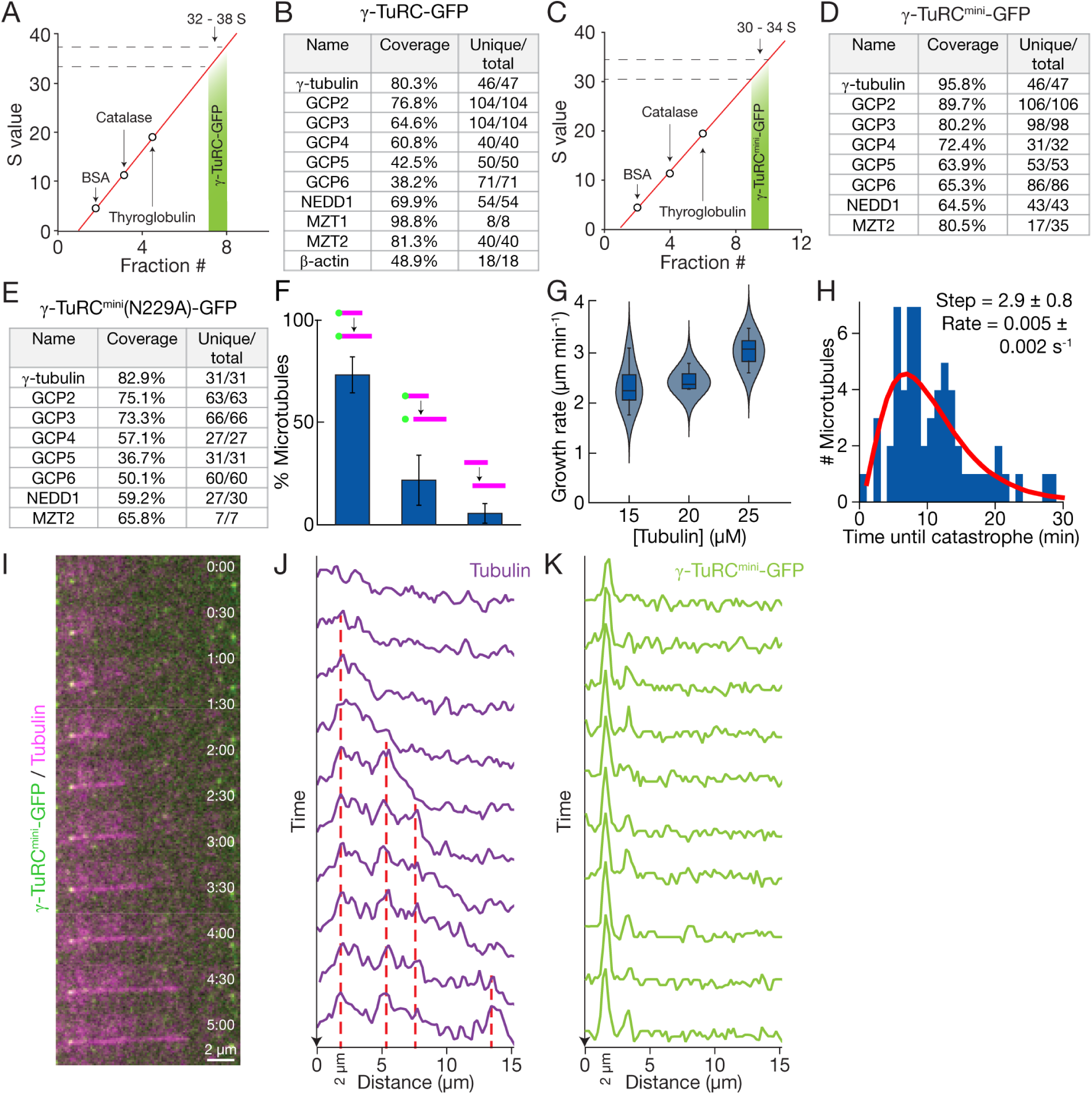
Biochemical characterization of reconstituted complexes and supplemental data for TIRF assays. A) Standard curve used to estimate S value of *γ*-TuRC-GFP from sucrose gradients. The red line is a linear fit to the peak sedimentation fractions for bovine serum albumin (BSA; 4.6 S), catalase (11.3 S), and thyroglobulin (19 S) identified in parallel sucrose gradients. Fractions 7 (∼32 S) and 8 (∼38 S) contained the highest concentration of *γ*-TuRC-GFP proteins and were pooled and used for subsequent biochemical and structural studies. B) Results from LC-MS/MS analysis of *γ*-TuRC-GFP. Coverage represents the % of the sequence that was identified by the LC-MS/MS analysis. Unique/total designates the number of unique peptides and total peptides respectively identified by the LC-MS/MS analysis. C) Standard curve used to estimate S value of *γ*-TuRC^mini^-GFP from sucrose gradients. The red line is a linear fit to the peak sedimentation fractions for bovine serum albumin (BSA; 4.6 S), catalase (11.3 S), and thyroglobulin (19 S) identified in parallel sucrose gradients. Fractions 9 (∼30 S) and 10 (∼34 S) contained the highest concentration of *γ*-TuRC^mini^-GFP proteins and were pooled and used for subsequent biochemical and structural studies. D) Results from LC-MS/MS analysis of *γ*-TuRC^mini^-GFP. E) Results from LC-MS/MS analysis of *γ*-TuRC^mini^(N229A)-GFP. F) The percentage of *γ*-TuRC^mini^-GFP-nucleated microtubules in a field of view that remained stably attached (left), temporarily attached (middle), or were not observed to be nucleated by (right) *γ*-TuRC^mini^-GFP (schematized above each bar). Data is presented as mean +/- s.e.m. n = 733 microtubule nucleation events, pooled from five independent experiments. G) Growth rates of microtubules nucleated by *γ*-TuRC^mini^-GFP at increasing tubulin concentrations. Solid lines drawn within box plots represent the mean; the boxes represent 25th and 75th percentile data; and the whiskers represent 5th and 95th percentile data. Violin plots show kernel density distributions of the data generated using a bandwidth value of 1 μm/min. n = 19 microtubule plus ends from two independent experiments were measured at each tubulin concentration. H) Distribution of lifetimes, or the “time until catastrophe”, of microtubules nucleated by *γ*-TuRC^mini^-GFP. The red line is a fit to the gamma distribution (Gardner et al., 2011), which yields a step size of 2.9 +/- 0.8 and a rate parameter of 0.005 +/- 0.002 s^-1^ (maximum likelihood estimates +/- 95% confidence intervals). n = 61 catastrophe events, pooled from five independent experiments. I) Two-color overlay time lapse sequence of a microtubule nucleated by *γ*-TuRC^mini^-GFP. J) Linescans of the growing microtubule in I) (tubulin channel) over each time frame. Dashed red lines indicate positions of persistent fluorescence intensity fluctuations (“speckles”) that do not appear to change in position over time. K) Linescans of the growing microtubule in I) (*γ*-TuRC^mini^-GFP channel) over each time frame. The *γ*-TuRC^mini^-GFP punctum (at ∼2.0 μm) co-localizes with the same position of the minus end of the nucleated microtubule in J).

**Figure S2.**
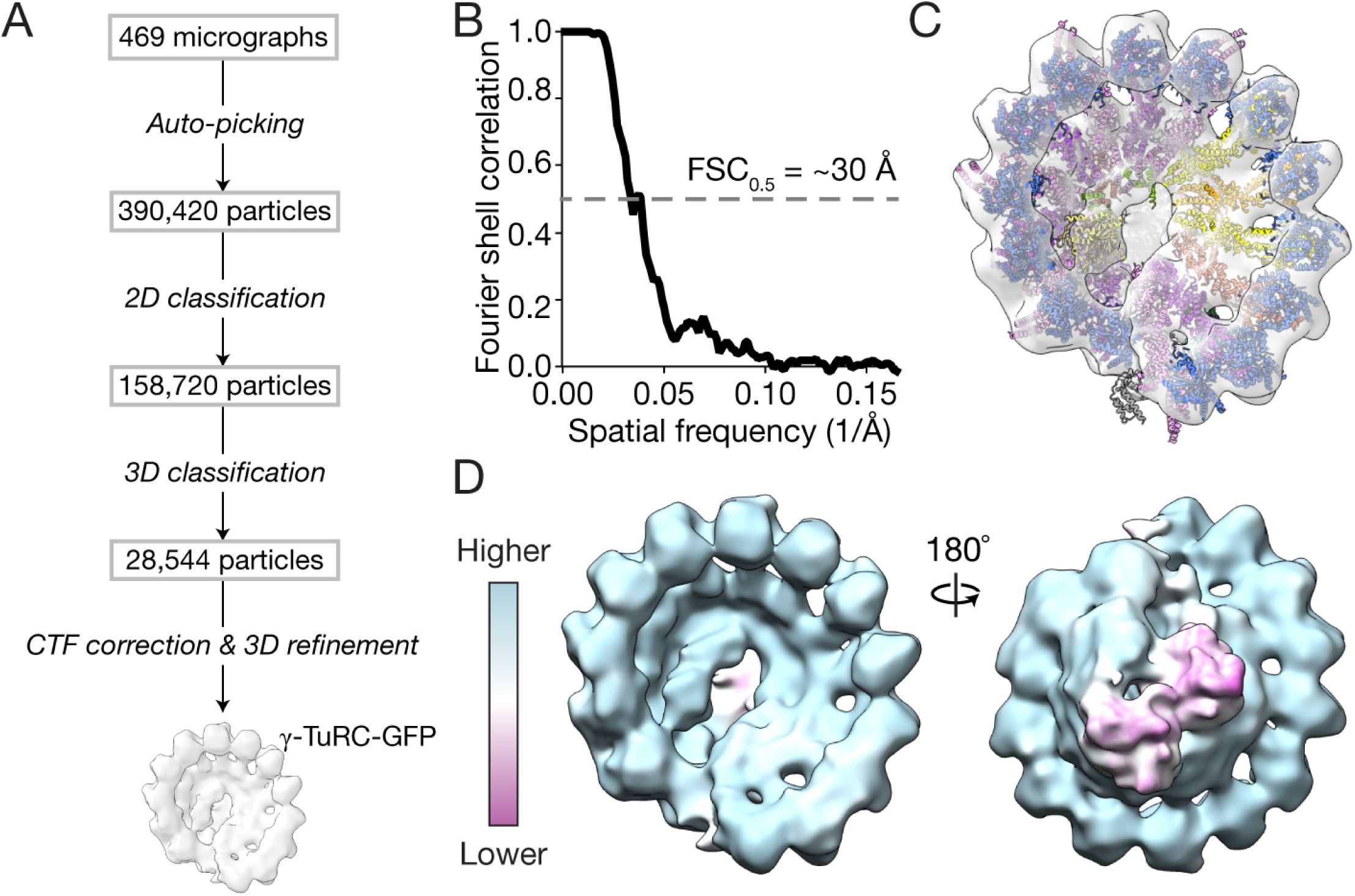
Supplemental data for negative stain EM analysis of *γ*-TuRC-GFP. A) Workflow for generating a negative stain EM 3D reconstruction of *γ*-TuRC-GFP. B) Masked FSC curve for the *γ*-TuRC-GFP reconstruction. FSC = 0.5 is indicated by a dashed grey line, and an estimate of the corresponding resolution is indicated. C) Rigid-body fit of the native human *γ*-TuRC model from Figure 1A into the *γ*-TuRC-GFP density map. D) Two views of the *γ*-TuRC-GFP density map colored according to local correlation of the model fit in C) (see Methods).

**Figure S3.**
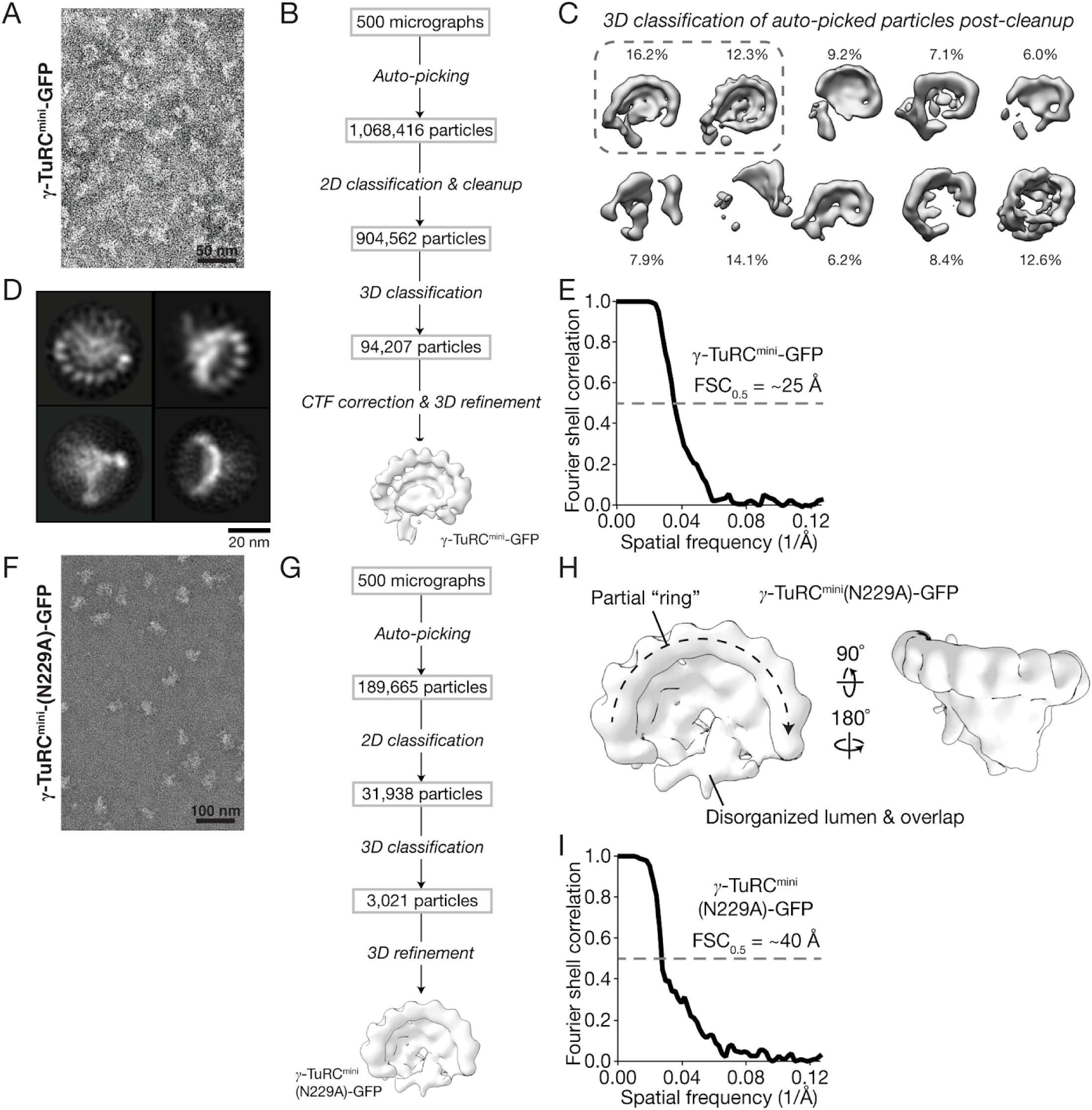
Supplemental data for negative stain EM analyses of *γ*-TuRC^mini^-GFP and *γ*-TuRC^mini^(N229A)-GFP. A) Transmission EM micrograph of negative stained *γ*-TuRC^mini^-GFP. B) Workflow for generating a negative stain EM 3D reconstruction of *γ*-TuRC^mini^-GFP. C) 3D classification results of *γ*-TuRC^mini^-GFP particles after initial cleanup. The percentage of total particles each class comprises is indicated. The dashed line encircles the two 3D classes used for downstream processing, which underwent a second round of 3D classification (classes not shown; this reduced the number of particles from ∼250,000 to 94,207 particles as shown in the workflow) prior to 3D refinement and final reconstruction of *γ*-TuRC^mini^-GFP. D) 2D class averages representing four different views of refined particles used to generate the *γ*-TuRC^mini^-GFP 3D reconstruction. E) Masked FSC curve for the *γ*-TuRC^mini^-GFP reconstruction. FSC = 0.5 is indicated by a dashed grey line, and an estimate of the corresponding resolution is indicated. F) Transmission EM micrograph of negatively-stained *γ*-TuRC^mini^(N229A)-GFP. G) Workflow for generating a negative stain EM 3D reconstruction of *γ*-TuRC^mini^(N229A)-GFP. H) Two views of a 3D reconstruction of *γ*-TuRC^mini^(N229A)-GFP from negative stain EM data. I) Masked FSC curve for the *γ*-TuRC^mini^(N229A)-GFP reconstruction. FSC = 0.5 is indicated by a dashed grey line, and an estimate of the corresponding resolution is indicated.

## Methods

### Expression and purification of *γ*-TuRC-GFP, *γ*-TuRC^mini^-GFP and *γ*-TuRC^mini^(N229A)-GFP from insect cells

Plasmids encoding Myc-His_6_-tagged human *γ*-tubulin, GCP2, GCP3, GCP4, GCP5, GCP6 (Murphy et al., 2001), GFP-tagged MZT2A (Teixidó-Travesa et al., 2010), and Myc-His_6_-tagged NEDD1 (Lüders et al., 2006) were kind gifts from Dr. Jens Luders. A plasmid containing the MZT1 coding sequence (pDONR223_C13orf37_WT) was a gift from Jesse Boehm, William Hahn & David Root (Addgene plasmid # 81872; http://n2t.net/addgene:81872; RRID:Addgene_81872; (Kim et al., 2016). A plasmid encoding for human *β*-actin was a kind gift from Dr. Gregory Alushin. Coding sequences were individually subcloned into the multiple cloning site of pACEBac1 (Bieniossek et al., 2008) using Gibson assembly (pACEBac1-MZT1 and pACEBac1-*β*-actin; (Gibson et al., 2009)) or standard restriction enzyme-based cloning methods (all other coding fragments). BstXI sites within coding sequences were removed by introducing silent mutations using primer-based site-directed mutagenesis. For *γ*-tubulin, a TEV protease site followed by a His_6_-tag and a stop codon were introduced at the 3’ end of the coding sequence. For MZT2, a ZZ-tag followed by a TEV cleavage site was PCR amplified from a dynein expression construct (Steinman et al., 2017) and inserted at the 5’ end of the MZT2 coding sequence. mEGFP with a 5’ PreScission protease site and a 3’ stop codon was PCR amplified from an in-house mammalian expression vector and inserted at the 3’ end of the MZT2 coding sequence.

Most *γ*-TuRC-GFP proteins could be individually overexpressed reasonably well, but GCP2 and GCP3 had very poor soluble expression levels. Studies of the *S. ceriviseae γ*-TuSC, comprised of Tub4 (*γ*-tubulin), Spc97 (GCP2), and Spc98 (GCP3), have shown that their purification yield is enhanced if co-overexpressed (Vinh et al., 2002). It seemed to us likely that a similar strategy could enhance the yield of not just GCP2 and GCP3, but potentially all 8 complex proteins. In initial tests, co-overexpressing all 10 proteins using a single baculovirus did increase soluble yields of GCP2 and GCP3, but only slightly. Further, we noticed that MZT2-mEGFP dominated protein expression levels, especially when codon optimized for expression in insect cells, but codon optimization of GCP2 and GCP3 had little effect on their expression levels. We therefore chose not to codon optimize any of the proteins in our final constructs. Finally, to further compensate for the poor expression of GCP2 and GCP3, we also generated a second bacmid containing only the human *γ*-TuSC proteins (*γ*-tubulin-TEV-His6, GCP2, and GCP3) and co-infected insect cells with both baculoviruses (Figure 1A), which we found significantly enhanced *γ*-TuRC-GFP yields. pACEBac1 vectors containing all 10 (pACEBac1-*γ*-TuRC-GFP) or just *γ*-tubulin-TEV-His6/GCP2/GCP3 (pACEBac1-*γ*-TuSC) were sequentially constructed using BstXI and I-CeuI restriction digestion and ligation, per the MultiBac system manual (Bieniossek et al., 2008). The final pACEBac1-*γ*-TuRC-GFP and pACEBac1-*γ*-TuSC plasmids were analyzed by asymmetric restriction digestion and complete Sanger sequencing of all open reading frames. pACEBac1-*γ*-TuRC^mini^-GFP was constructed in a similar manner. To construct *γ*-TuRC^mini^(N229A)-GFP, an N229A point mutation was introduced into the pACEBac1-*γ*-tubulin-TEV-His6 plasmid using primer-based site directed mutagenesis. pACEBac1-*γ*-TuRC^mini^(N229A)-GFP and pACEBac1-*γ*-TuSC(N229A) donor plasmids were sequentially constructed using BstXI and I-CeuI restriction digestion and ligation, as for pACEBac1-*γ*-TuRC^mini^-GFP and pACEBac1-*γ*-TuSC.

Polycistronic donor plasmids were transformed into DH10MultiBacTurbo cells (ATG:biosynthetics GmbH) and transposition-positive colonies were selected and used to generate recombinant bacmids. Bacmids were transfected into Sf9 cells per the Bac-to-Bac manual (Invitrogen), baculoviruses were amplified twice, and fresh P3 virus from *γ*-TuRC-GFP/ bacmid and *γ*-TuSC bacmid were mixed together at a 1:1 ratio. This virus mixture was used to infect 1.8 - 2.4 L of High Five cells at a cell density of 3 x 10^6^/mL for 60 hr at 27 °C. Cells were harvested by centrifugation at 1000 x g, resuspended in 80 mL ice-cold lysis buffer (40 mM HEPES pH 7.5, 150 mM KCl, 1 mM MgCl2, 10% glycerol (v/v), 0.1% Tween-20, 0.1 mM ATP, 0.1 mM GTP, 1 mM 2-mercaptoethanol, 4 cOmplete EDTA-free Protease Inhibitor Cocktail tablets (Roche), 500 U benzonase, 2 mM PMSF, and 4 mM benzamidine-HCl), and lysed by dounce homogenization on ice. The lysate was clarified at 322,000 x g for 1 hr at 4 °C, 0.22 µm syringe-filtered, and loaded onto a 1 mL NHStrap column (GE Healthcare) previously coupled to 10-20 mg rabbit IgG (Innovative Biosciences) following the manufacturer’s instructions. The IgG column was washed with lysis buffer followed by gel filtration buffer (40 mM HEPES pH 7.5, 150 mM KCl, 1 mM MgCl2, 10% glycerol (v/v), 0.1 mM GTP, and 1 mM 2-mercaptoethanol). 1 mg of TEV protease (stored in 40 mM HEPES pH 7.5, 30% (w/v) glycerol, 150 mM KCl, 1 mM MgCl2, and 3 mM 2-mercaptoethanol) was diluted into 1 mL of gel filtration buffer, injected onto the column, and proteolysis was allowed to proceed for 2 hr at 4 °C. The digested eluate was pooled, concentrated by dialyzing against dialysis buffer (40 mM HEPES pH 7.5, 150 mM KCl, 1 mM MgCl2, 60% sucrose (w/v), 0.1 mM GTP, and 2 mM 2-mercaptoethanol) for 4 hr at °C, and gel filtered over a Superose 6 10/300 GL or a Superose 6 Increase 10/300 GL column pre-equilibrated in gel filtration buffer. The peak fractions were identified by SDS-PAGE followed by Coomassie staining and/or negative stain TEM, pooled and loaded onto a 2 mL sucrose gradient composed of 10%, 20%, 30% and 40% sucrose (w/v) in gradient buffer (40 mM HEPES pH 7.5, 150 mM KCl, 1 mM MgCl2, 0.01% Tween-20 (v/v), 0.1 mM GTP, and 1 mM 2-mercaptoethanol). The gradient was centrifuged at 50,000 rpm in a TLS-55 rotor at 4 °C for 3 hr with minimum acceleration and no break. Fractions were manually collected with a cut-off P1000 pipette tip and analyzed by SDS-PAGE followed by Coomassie staining and/or negative stain TEM. Peak fractions were aliquoted, snap frozen and stored in liquid N_2_. For estimating sedimentation coefficients, parallel gradients were run with 200 µg each of BSA, catalase, and thyroglobulin (GE Healthcare). Gradients were fractionated into 250 µL (Figure S1A) or 150 µL (Figure S1C) and analyzed by SDS-PAGE followed by Coomassie staining.

### Mass spectrometry

A 40 μL aliquot of purified recombinant complex was thawed and mixed with 10 μL of 5X sample buffer (Tris-HCl, pH 6.8, 10% (w/v) SDS, 50% (w/v) glycerol, 700 mM 2-mercaptoethanol, and 0.25% (w/v) bromophenol blue). After boiling, the sample was loaded into a single lane of a 4%–20% Tris-glycine pre-cast gel with “wide wells” (Novex) and allowed to migrate ∼1 cm into the stacking gel. A corresponding ∼1 cm x 1 cm gel plug was cut out of the gel, further cut into 1 mm cubes. Protein gel bands were destained, reduced, alkylated, and digested with Endopeptidase Lys-C and trypsin overnight. Peptides were extracted with 70% acetonitrile in 5% formic acid at 35 °C, dried and redissolved in 10uL. 3uL was injected and separated using a gradient increasing from 2% B to 35% B over 70 min (Buffer A: 0.1% formic acid; Buffer B: 80% acetonitrile in 0.1% formic acid). Peptides were resolved using a 12 cm 75 µm column with 3 µm C18 particles (Nikkyo Technos Co., Ltd. Japan) coupled to a Fusion Lumos mass spectrometer (Thermo Scientific) operated in high/high mode. Raw data were analyzed with MaxQuant 1.6.2.10.

### Quantitative western blotting

Complex concentrations were estimated using quantitative western blotting against *γ*-tubulin standards using anti-*γ*-tubulin (clone GTU-88; Sigma), assuming that each complex contains 14 *γ*-tubulin molecules. *γ*-tubulin was expressed in insect cells using the pACEBac1-*γ*-tubulin-TEV-His6 plasmid and purified as previously described (Rice et al., 2008).

### Purification and biotinylation of GFP nanobody

The construct for a recombinant GFP nanobody (clone LaG-16; (Fridy et al., 2014)) was a kind gift from Dr. Michael Rout. The plasmid was transformed into Rosetta2(DE3) cells and periplasmic overexpression was induced with 0.1 mM IPTG for 16 hours at 16 °C. Cells from 6 L of culture were harvested at 5,000 X g for 10 min and resuspended in TES buffer (200 mM Tris-HCl, pH 8.0, 0.5 mM EDTA, 500 mM sucrose). The cells were osmotically shocked on ice for 30 min by diluting them 5-fold in 1:4 water:TES buffer. Periplasmic extract was separated from cell debris by centrifugation for 10 min at 6,000 x g, and this supernatant was further clarified at 20,000 x g for 20 min at 4 °C. The final supernatant was supplemented with NaCl to 150 mM and incubated with 3 mL His60 resin (Takara Biosciences) for 30 min. The resin was washed in batch with wash buffer 1 (20 mM Tris-HCl, pH 8.0 and 900 mM NaCl), wash buffer 2 (20 mM Tris-HCl, pH 8.0, 150 mM NaCl, and 10 mM imidazole), loaded onto a disposable column, and the protein was eluted with Ni-NTA elution buffer (20 mM Tris-HCl, pH 8.0, 150 mM NaCl, and 250 mM imidazole). Peak fractions were identified by Bradford assay, pooled, and concentrated with a 10 kDa cutoff spin filter (Millipore). To produce biotinylated GFP nanobody, NHS-biotin (Sigma) was dissolved in DMSO and added to the concentrated protein at a molar ratio of 10:1 such that the final DMSO concentration did not exceed 5%. The reaction was incubated at 4 °C for 4 hours and clarified by centrifugation at 21,000 x g for 20 min at 4 °C. The nanobody was then gel filtered over a Superdex 75 10/300 column equilibrated in coupling buffer (150 mM sodium bicarbonate, pH 8.0, 150 mM NaCl). Biotinylated antibody was supplemented with glycerol to 10% (v/v), flash frozen in liquid N2 and stored at −80 °C.

### Negative stain electron microscopy and data processing

3 µL of sucrose density gradient-purified *γ*-TuRC-GFP, *γ*-TuRC^mini^-GFP, or *γ*-TuRC^mini^(N229A)-GFP was applied to glow discharged carbon-coated copper grids (CF-400-Cu; EMS) and incubated for 45 s on ice. Protein solution was removed by manual blotting with Whatman No. 1 filter paper, then 3 µL was applied again to improve surface density. This procedure was optionally repeated resulting in a total of 2-4 applications, depending on the concentration of *γ*-TuRC complex. After the final application, protein was manually blotted from one side of the grid while freshly filtered 1 or 2% uranyl acetate (w/v) was simultaneously pipetted from the opposite side to exchange the solution. Grids were incubated in uranyl acetate for a further 45 s. Stain was removed by manual blotting and grids were air dried for >24-48 hr in a sealed container containing desiccant prior to imaging. TEM micrographs were recorded on an FEI Tecnai G2 microscope operating at 120 kV at a magnification of 30,000X (3.96 Å/pixel) or 36,000X (3.036 Å/pixel) using a BioSprint 29 CCD camera.

Images were cropped in Fiji to remove micrograph metadata labels (Schindelin et al., 2012), and .tif formats were converted to .mrc using RELION’s “relion_image_handler” function. CTF parameters were estimated using CTFFIND4 (Rohou and Grigorieff, 2015). All subsequent processing was done in RELION version 3.1 and UCSF Chimera (Zivanov et al., 2018; Pettersen et al., 2004). The main 3D reconstruction workflow steps for *γ*-TuRC-GFP, *γ*-TuRC^mini^-GFP, or *γ*-TuRC^mini^(N229A)-GFP are outlined in Figures S2A, S3B, and S3G, respectively. Generally, micrographs were imported and a small set of <1,000 particles was manually picked and subjected to reference-free 2D classification. The resulting set of averages were used as initial templates for 1-2 rounds of auto-picking using RELION’s built-in implementation yielding best templates for optimal picking. Auto-picked particles were binned by 2 and subjected to reference-free 2D classification to remove particles likely corresponding to dirt and other contaminants. A random subset of the cleaned, binned particles was used to generate an *ab initio* model. Then, all cleaned, binned particles were subjected to an intermediate 3D auto-refinement step using the *ab initio* model as a reference, re-extracted using the refined co-ordinates, and subjected to 3D classification. Particles that generated 3D classes containing the highest level of detail were re-extracted from CTF-corrected micrographs at the unbinned pixel size (3.036 Å or 3.96 Å) and subjected to a final round of 3D auto-refinement using one of the classes as a reference model.

A composite model of the native human *γ*-TuRC was constructed by substituting our recently published models for the lumenal bridge and MZT2/GCP2-NHD/*γ*-TuNA (PDB ID: 6×0U and 6×0V; (Wieczorek et al., 2020)) with the corresponding poly-alanine models for the lumenal bridge, the HB, and the CC regions in our previous structure of the native complex (PDB ID: 6V6S; (Wieczorek et al., 2019)). A poly-alanine model for the unassigned MZT “module” was generated with MZT1/GCP6-NHD and rigid-body fitted into the unassigned MZT module density found at the overlap region, as reported in (Wieczorek et al., 2020). The composite model was then rigid-body fitted into density maps using the “Fit in map” function of Chimera. The local correlation between the model and the density map after fitting was calculated using Chimera’s “vop localCorrelation” command with a window size of 5 voxels. All figures displaying density maps and protein models were generated using UCSF Chimera (Pettersen et al., 2004) or UCSF ChimeraX (Goddard et al., 2018).

### Turbidity-based microtubule nucleation assay

For all in vitro assays used in this study, tubulin purified from calf brain (Gell et al., 2011) was quickly thawed at 37 °C and centrifuged at 350,000 x g for 10 min at 4 °C. The concentration of the supernatant was measured spectrophotometrically using an extinction coefficient of 115,000 M^-1^ cm^-1^ and a molecular weight of 110 kDa for the tubulin dimer. Tubulin prepared in this way was kept on ice for no more than 2 hr.

Microtubule nucleation was assayed by preparing 75 µL reactions containing 0.25 nM *γ*-TuRC-GFP, *γ*-TuRC^mini^-GFP, or *γ*-TuRC^mini^(N229A)-GFP, 20 µM tubulin, and 1 mM GTP in BRB80 (80 mM K-PIPES, pH 6.8, 1 mM MgCl2, and 1 mM EGTA). An appropriate volume of 30% sucrose gradient buffer (see *γ*-TuRC-GFP purification) was included for tubulin-only controls. Components were mixed and incubated on ice for 5 min. Reactions were split and loaded into a pre-warmed 384-well, clear-bottomed microplate in duplicate (∼32 µL reaction per well), and 15 µL of fluorescence-free immersion oil (Cat 16212, Cargille Laboratories) was layered over each reaction. This loading time was limited to a total of 5 minutes. The microplate was immediately placed on a BioTek microplate reader warmed to 37 °C and absorbance measurements at 350 nm were initiated at a frequency of 1 per minute for a total of 1 hour.

### Single molecule *γ*-TuRC-GFP microtubule nucleation assay

The TIRF microscope setup was as described in (Ti et al., 2018), except that a 1.5 X optovar was introduced in the optical path and images were acquired on a Photometrics Prime 95B sCMOS camera. Tubulin purified from calf brains was functionalized with X-rhodamine succinimidyl ester following established protocols (Hyman et al., 1991). Glass coverslips (Gold Seal 18 x 18 mm, No. 1) and glass slides (Buehler 40-80000-01) were rinsed in acetone, sonicated for 20 min in 50% (v/v) methanol, sonicated for 20 min in 0.5 M KOH, and rinsed with milliQ water. After drying with nitrogen gas, the coverslips were plasma cleaned for 5 min (Harrick Plasma PDC-32G) and adhered to glass slides via two strips of double-sided tape separated by ∼5 mm. The flow cell was rinsed with BRB80 + 1 mM TCEP, followed by 0.2 mg/mL PLL-PEG-biotin (PLL(20)-g[3.5]-PEG(2)/PEG(3.4)-biotin(20%); SUSOS AG) prepared in BRB80 + 1 mM TCEP. After 5 min, the flow cell was rinsed with BRB80 + 1 mM TCEP, and a mixture containing 0.5 mg/mL κ-casein and 0.25 mg/mL neutravidin prepared in BRB80 + 1 mM TCEP was flowed in. After 5 min, the flow cell was rinsed with assay buffer (BRB80 + 1 mM TCEP, 50 mM KCl, 0.15% (w/v) methylcellulose, 0.2 mg/mL κ-casein, and 1 mM GTP), and 0.02 mg/mL biotinylated GFP nanobody prepared in assay buffer was flowed in. After 5 min, the flow cell was rinsed with assay buffer and ∼1 pM *γ*-TuRC-GFP prepared in assay buffer was flowed in. After 5 min, unbound *γ*-TuRCs were rinsed out with assay buffer and a reaction mixture containing 15 µM tubulin (with ∼5% X-rhodamine-labeled tubulin) and oxygen scavengers (0.035 mg/mL catalase, 0.2 mg/mL glucose oxidase, 2.5 mM glucose, and 10 mM DTT) was prepared in assay buffer and introduced to the flow cell. The flow cell was sealed with VALAP and placed on the TIRF microscope stage. Two-color images were acquired at 10 s per frame and an exposure time of 500 ms (tubulin channel) or 100 ms (GFP channel). Total imaging times were between 20 - 30 min. The microscope chamber was heated to ∼35 °C prior to image acquisition.

Time lapse images were drift-corrected using the MultiStackReg plugin in Fiji made by Brad Busse. The number of *γ*-TuRC^mini^-GFP puncta was measured using the “Spot Intensity Analysis” Fiji plugin built by Nico Stuurman and Ankur Jain. *γ*-TuRC^mini^-GFP-mediated microtubule nucleation events were counted and tracked manually in Fiji. Only nucleation events which initiated from *γ*-TuRC^mini^-GFP puncta were considered in the analysis in Figure 3D-E. Microtubule growth rates were calculated by estimating the change in length of microtubules growing from *γ*-TuRC^mini^-GFP puncta divided by the length of time spent in the growth phase. Growth rates of microtubules that dissociated from GFP puncta were only measured for the time that they were associated with *γ*-TuRC^mini^-GFP. Figure 4E shows the classification of microtubule nucleation events and whether or not they were attached to *γ*-TuRC-GFP. The length of time that *γ*-TuRC^mini^-GFP-associated microtubules spent in the growth phase before undergoing catastrophe was measured and is plotted in Figure S1H. A gamma distribution was used to fit these data, as previously described (Gardner et al., 2011), which produced similar values for the rate (0.005 +/- 0.002 s^-1^) and step (2.9 +/- 0.8) parameters as for microtubules nucleated from GMPCPP seeds. Fitting was performed in MATLAB (MathWorks) and SciDAVis (http://scidavis.sourceforge.net/).

## Data availability

The data that support the findings of this study are available from the corresponding author upon reasonable request.

## Notes

### Competing Interest Statement

The authors have declared no competing interest.

